# Pre-Existing Serotonergic Pathways Guide the Navigation of Developing Serotonergic Axons

**DOI:** 10.64898/2026.09.04.749432

**Authors:** Marta Picchi, Sara Migliarini, Serena Nazzi, Giulia Gianni, Skirmantas Janušonis, Noemi Barsotti, Massimo Pasqualetti

## Abstract

The serotonergic system originates from a small population of brainstem neurons whose axons form one of the most extensive projection networks in the vertebrate nervous system. Although a number of molecular regulators of serotonergic development have been identified, the core principles that organize this widespread axonal architecture remain poorly understood. Classical neuroanatomical studies have proposed that serotonergic axons may navigate by growing along pre-existing fiber tracts, a process termed epiphytic guidance, but this hypothesis has remained largely untested.

Here, we used organotypic transplantation assays to investigate how embryonic serotonergic axons navigate within developing and adult neural tissue. Rostral raphe explants from Tph2-GFP embryos were grafted onto embryonic hindbrain flat-mounts or adult brain slices, allowing donor-derived axons to be visualized in relation to genetically labelled endogenous serotonergic pathways and host tissue architecture. Across different grafting configurations, developmental stages, and both homotopic and heterotopic host territories, donor-derived axons consistently aligned with pre-existing serotonergic pathways adopting local trajectories rather than growing independently of them. This substrate-dependent behavior extended to adult tissue, where transplanted embryonic axons preferentially followed pre-existing serotonergic axon directions. These observations reveal substantial navigational plasticity and indicate that axonal and tissue architectures can provide permissive and potentially instructive substrates for the dispersal of serotonergic fibers. Together, our findings provide experimental support for epiphytic guidance and suggest that serotonergic pathway assembly relies, at least in part, on a pioneer-follower mechanism of progressive self-scaffolding. Such a strategy may help explain how a small population of raphe neurons generates an extensive and spatially coherent neuromodulatory system.

## Introduction

The serotonin (5-HT) system constitutes one of the most widely distributed neuromodulatory networks of the vertebrate central nervous system^1,2^. A relatively small population of neurons located in the brainstem raphe nuclei gives rise to extensive ascending and descending axonal projections that innervate virtually all regions of the brain and spinal cord^3,4^. Unlike point-to-point projection systems, the serotonergic network ultimately forms a diffuse axonal fiber meshwork that permeates large territories of neural tissue, thereby allowing serotonin to influence sensory processing, emotional behavior, cognition, sleep-wake regulation, autonomic functions, and motor control^5–8^. In addition to these well-established functions in the mature nervous system, serotonin also acts as an important developmental signal, regulating processes such as neurogenesis, neuronal differentiation, migration, axonal branching, and circuit formation^9–11^.

The anatomical organization of the serotonergic system raises a fundamental developmental question: how is such a globally distributed axonal architecture assembled during embryogenesis? Serotonergic neurons are among the earliest neuronal populations to differentiate in the developing brainstem, and their axons rapidly extend over long distances to invade the forebrain, the hindbrain, and spinal territories^1,2,12^. Although the population-level distribution of serotonergic fibers has been extensively described in classical neuroanatomical studies, the cellular principles that organize their trajectories within developing tissue remain poorly understood. Several transcriptional regulators and molecular effectors have been implicated in serotonergic neuron specification, axonal growth and arborization, including Lmx1b, Pet-1/Fev, GAP-43, BDNF, protocadherin-α family members and other cell-adhesion or guidance-related molecules^13–20^.

These studies indicate that serotonergic axon development depends on both intrinsic genetic programs and extrinsic tissue-derived signals. However, they do not fully explain how a sparse population of early-born neurons generates an extensive and spatially coherent projection system across the entire neuraxis.

A general principle emerging from studies of developing nervous systems is that early-growing axons can profoundly influence the behavior of later-growing axons. In many systems, pioneer axons establish initial trajectories that are subsequently used by follower axons, thereby creating a structural scaffold for the progressive assembly of neural pathways^21^. Experimental evidence from invertebrate and vertebrate model organisms, including *Drosophila*, zebrafish, and *C. elegans*, has consistently shown that pioneer axons do not simply reach their targets independently, but progressively generate physical and molecular information that follower axons exploit for navigation, increasing the robustness and efficiency of circuit assembly^22–24^. This pioneer-follower organization thus provides an efficient developmental strategy through which complex neuronal networks can emerge from a limited set of initial guidance events.

Whether a similar principle contributes to the formation of the serotonergic projection system remains largely unresolved. Classical developmental studies have noted that serotonergic axons frequently grow in close association with pre-existing fiber tracts, including pathways that are established before or during the earliest phases of serotonergic innervation^1–3^. These observations have led to the proposal that serotonergic axons may use other axonal bundles as physical substrates for elongation, a process referred to as epiphytic guidance by analogy with non-parasitic plants that grow on other plants for physical support. In this view, serotonergic axons are not guided exclusively by diffusible chemotropic cues but also navigate by maintaining direct contact with pre-existing cellular architectures. Despite its conceptual relevance, this hypothesis has remained difficult to test experimentally, largely because it requires preserving the three-dimensional organization of neural tissue while visualizing the behavior of developing serotonergic axons in relation to endogenous fiber systems

An even more intriguing possibility is that this principle may operate within the serotonergic system itself. The first serotonergic axons to enter a territory may not simply represent the earliest output of a genetically specified developmental program; they may also generate part of the structural information required for subsequent serotonergic growth. Under this scenario, later-growing serotonergic axons would behave, at least in part, as follower axons, using pre-existing serotonergic fibers as permissive substrates for elongation and directional organization. The developing serotonergic system would therefore assemble through a self-scaffolding process in which the emerging network progressively contributes to its own expansion.

Organotypic transplantation and co-culture approaches provide a useful experimental framework for examining how developing axons respond to local tissue environments while preserving much of the native tissue architecture^25–27^. Building on this approach, we investigated this possibility by using an organotypic transplantation paradigm designed to test how developing serotonergic axons respond to different tissue environments. Embryonic raphe explants from Tph2-GFP mice^28^ were grafted onto embryonic hindbrain preparations or onto adult brain slices preserving endogenous cytoarchitecture and pre-existing axonal networks. This approach allowed us to examine whether transplanted serotonergic axons grow independently of the host tissue or whether their trajectories become organized by local axonal substrates. We show that developing serotonergic axons display distinct growth behaviors depending on the host environment and progressively align with endogenous serotonergic pathways after contact. These findings support the idea that pre-existing serotonergic fibers can act as permissive structural substrates for later-growing serotonergic axons, suggesting that the assembly of the serotonergic projection system may rely, at least in part, on a pioneer-follower-like mechanism of self-generated axonal scaffolding.

## Results and Discussion

### Embryonic hindbrain flat-mounts retain serotonergic viability and developmental progression *ex vivo*

A major challenge in studying serotonergic axon navigation is maintaining the three-dimensional organization of the developing neural tissue while preserving experimental accessibility. To address this, we established an embryonic hindbrain flat-mount organotypic preparation that retains the overall cytoarchitecture of the developing hindbrain and allows direct imaging of endogenous serotonergic axon growth over time.

To assess whether this preparation supports continued serotonergic development *ex vivo*, E12.5 hindbrain flat-mounts were either fixed immediately after dissection or maintained *in vitro* for three days, a duration empirically determined to provide the optimal balance between developmental progression and tissue quality. Flat-mounts were then processed for serotonin (5-HT) immunohistochemistry. Freshly dissected preparations displayed serotonergic neurons confined to two ventral longitudinal clusters within the raphe region and only sparse rostrally-directed axons (Fig. 1A). After three days in culture, serotonergic neurons showed evidence of lateral dispersal from the initial clusters. Concurrently, rostrally projecting serotonergic fibers had become considerably more numerous and extended further along the ascending pathway, consistent with ongoing axonal growth *ex vivo* (Fig. 1B).

**Figure 1.**
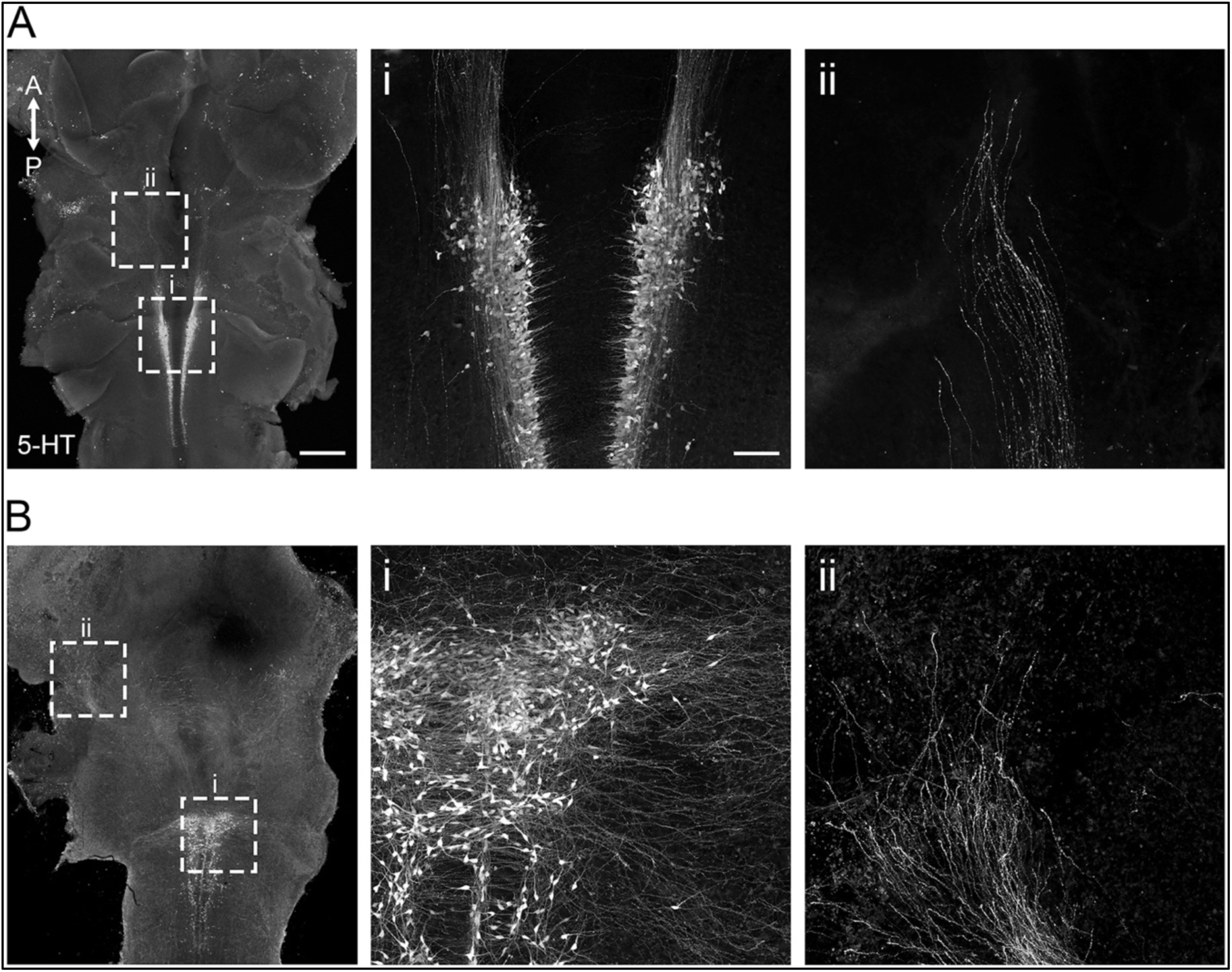
Embryonic hindbrain flat-mounts support continued serotonergic development *ex vivo*. **(A)** 5-HT immunofluorescence of an E12.5 hindbrain flat-mount fixed upon dissection. Endogenous serotonergic neurons are organized in two ventral longitudinal clusters and only a limited number of fibers extend rostrally. Dashed boxes indicate the regions shown at higher magnification in panels i and ii. **(i)** Higher-magnification view of the bilateral serotonergic cell clusters and **(ii)** of the most rostral serotonergic fibers present at the time of dissection. **(B)** 5-HT immunofluorescence of an E12.5 hindbrain flat-mount maintained *in vitro* for 3 days. Dashed boxes indicate the regions shown at higher magnification in panels i and ii. **(i)** Serotonergic neurons display marked lateral dispersal from the initial longitudinal clusters and are associated with an expanded local axonal network. **(ii)** Rostral serotonergic fibers are more numerous and extend along the ascending pathway. Anterior-posterior (A-P) orientation is indicated. 5-HT, serotonin. Scale bars: A and B, 500 μm; Ai-ii and Bi-ii, 100 μm.

These observations indicate that the flat-mount preparation supports continued serotonergic development while preserving the spatial organization of the developing hindbrain, providing an appropriate experimental substrate for investigating how transplanted serotonergic axons navigate within an intact developmental environment.

### Embryonic raphe explants engraft successfully and extend serotonergic axons into the host environment

To investigate how serotonergic axons navigate within developing neural tissue, we established an organotypic grafting paradigm based on embryonic flat-mount brain preparations using Tph2-GFP embryos as a source of donor 5-HT neurons. Explants were obtained under a fluorescence stereomicroscope by dissecting the rostral portion of the GFP-positive hindbrain domains, corresponding approximately to rhombomeres 1 and 2, with a medial cut that isolated half of the rhombomeric segment. The explant was then carefully trimmed under fluorescence guidance to remove as much non-fluorescent tissue as possible, in order to limit the graft to the extent of the 5-HT neuron cluster (Fig. 2A). This procedure yielded a well-defined donor fragment containing the earliest-born serotonergic neurons of the dorsal raphe anlage, identifiable by their intense GFP fluorescence.

**Figure 2.**
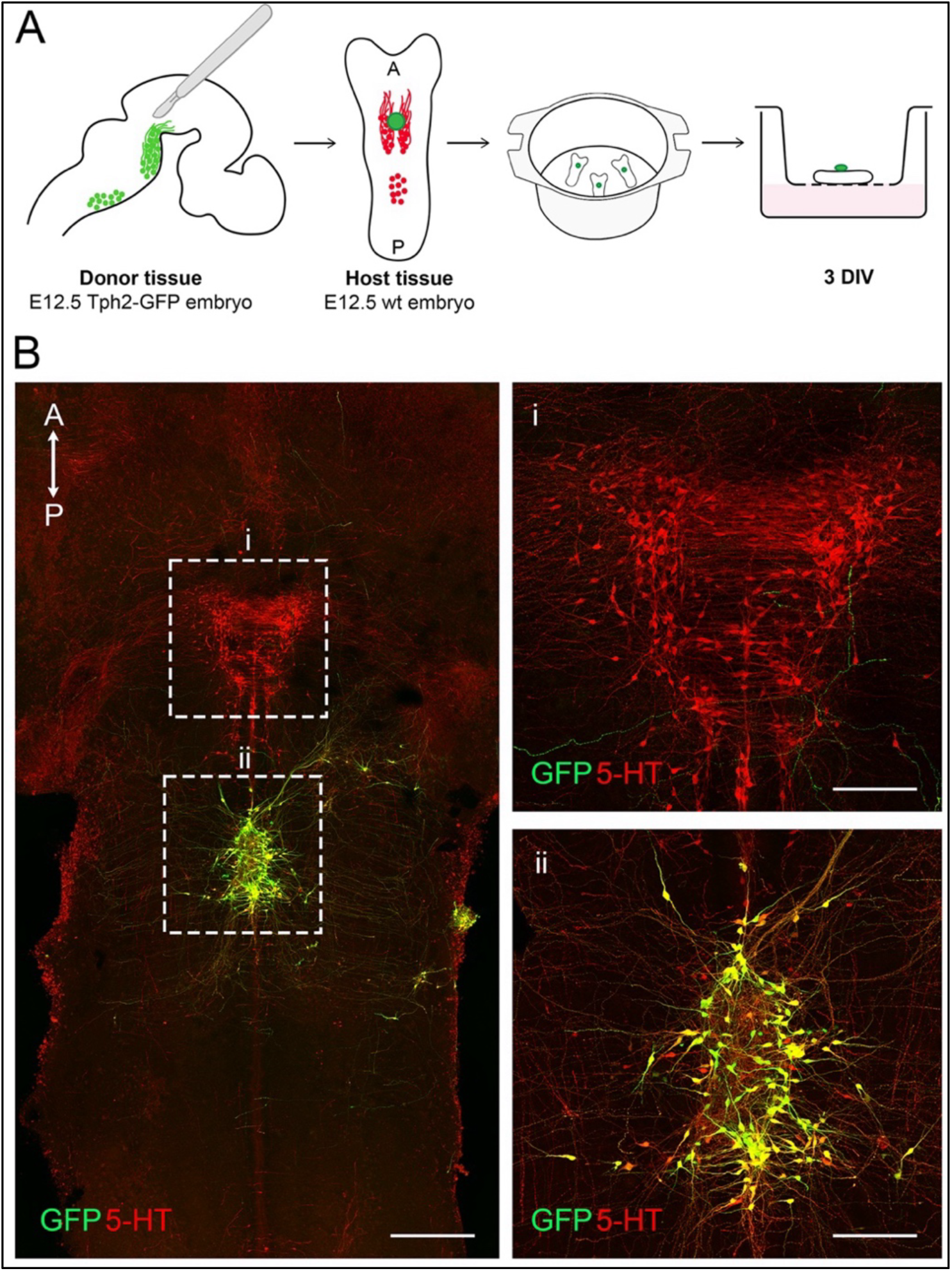
Embryonic serotonergic neurons retain their identity and extend axons into the host hindbrain. **(A)** Schematic overview of the experimental design used for embryonic transplantation. Donor tissue containing rostral raphe serotonergic neurons was dissected from the ventral hindbrain of an E12.5 Tph2-GFP embryo and homochronically grafted onto the ventricular surface of a wild-type hindbrain flat-mount. Preparations were maintained 3 days *in vitro* on membrane inserts. **(B)** Whole-mount merged image showing donor-derived GFP-positive cells and fibers in green and 5-HT immunoreactivity in red. The graft remained viable and extended numerous GFP-positive axons into the surrounding host tissue at 3 DIV. Dashed boxes indicate the regions shown at higher magnification in panels i and ii. **(i)** High magnification of the endogenous host serotonergic domain, showing 5-HT-positive neurons and fibers together with donor-derived GFP-positive axons extending into the surrounding tissue. **(ii)** High magnification of the graft and adjacent axons. GFP-positive donor neurons show extensive colocalization with the 5-HT immunosignal (yellow), confirming the preservation of their serotonergic identity. Anterior-posterior (A-P) orientation is indicated. 5-HT, serotonin; DIV, days *in vitro*. Scale bars: B, 500 μm; Bi-ii, 200 μm.

Host flat-mount preparations derived from wild-type embryos were positioned with the pial surface facing the membrane insert and the ventricular surface exposed upward, allowing donor explants to be placed directly onto the ventricular side of the neural tube. Following three days in culture, GFP-positive neurons of the graft remained fluorescent and viable, with some cells showing limited displacement from the donor tissue, consistent with early migratory behavior (Fig. 2B). Most strikingly, donor neurons extended large numbers of GFP-positive axons that spread extensively into the surrounding host tissue. Double immunostaining for GFP and 5-HT confirmed that these axons were serotonergic, demonstrating that donor neurons maintained their neurochemical identity throughout the culture period (see panel ii in Fig. 2B).

These observations establish that the organotypic grafting paradigm supports the survival, phenotypic maintenance, and axonal growth of transplanted serotonergic neurons, providing a suitable experimental framework to examine how donor-derived serotonergic axons interact with the host tissue environment.

### Serotonergic axons progressively acquire rostro-directed orientation within the host tissue

To determine whether donor-derived serotonergic axons exhibit preferential patterns of navigation within the developing hindbrain, we first examined their overall distribution throughout the host tissue. Whole-mount maximum projections showed that GFP-positive fibers spread broadly around the graft and appeared to lack a clear directional preference. However, this apparent lack of directionality resulted largely from combining axons located at different tissue depths into a single maximum projection (Fig. 3A). We therefore analyzed successive groups of confocal optical sections independently by measuring the relative optical density of the GFP-positive signal within rostral, caudal, and lateral quadrants surrounding the graft (Fig. 3B).

**Figure 3.**
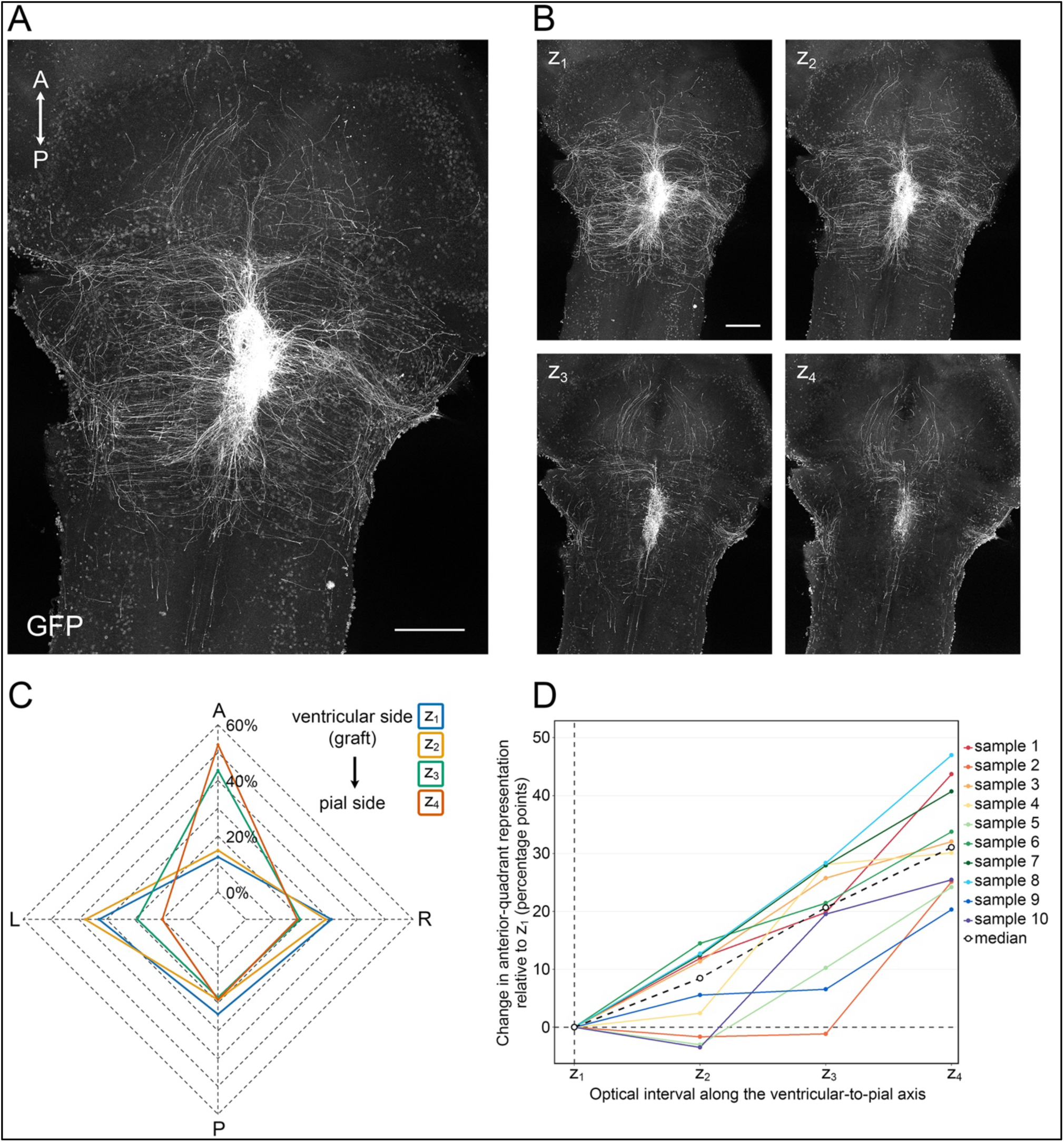
Donor-derived serotonergic axons show a depth-dependent transition from radial to rostro-directed growth. **(A)** Maximum-intensity projection of GFP-positive axons extending from an E12.5 Tph2-GFP rostral raphe explant homochronically and homotopically grafted onto the ventricular surface of a wild-type hindbrain flat-mount and maintained 3 days *in vitro*. Projection of the complete z-stack shows donor-derived fibers extending broadly around the graft. Anterior-posterior (A-P) orientation is indicated. **(B)** Maximum-intensity projections of four successive optical intervals, z_1_-z_4_, each comprising 10 consecutive confocal sections. z_1_ is closest to the ventricular surface and the graft, whereas z_4_ is closest to the pial surface (z_1_ = 0-18 μm, z_2_ = 20-38 μm, z_3_ = 40-58 μm, z_4_ = 60-78 μm). Donor-derived fibers are broadly distributed in the most ventricular intervals and become progressively enriched in the anterior quadrant closer to the pia. **(C)** Radar plot showing the percentage distribution of GFP relative optical density of the anterior, posterior, left, and right quadrants for each of the four optical intervals shown in B. **(D)** Depth-dependent change in the percentage of GFP signal represented within the anterior quadrant for all analyzed explants. For each preparation, the value measured in the most ventricular interval (z1) was used as an internal reference and subtracted from the values measured at all subsequent optical intervals; z1 was therefore set to 0, representing no change from baseline. Values are expressed as percentage-point differences relative to z1. Colored lines represent individual explants, and the black dashed line and open circles indicate the median. Statistical analysis of the original, non-normalized percentages revealed a significant overall effect of optical depth on rostral axon representation (Friedman test, χ²(3) = 25.56, *p* < 0.0001; *n* = 10 explants). A, anterior; P, posterior; L, left; R, right. Scale bars: 500 μm.

This analysis revealed a clear depth-dependent reorganization of donor-derived axonal trajectories. In the optical sections closest to the ventricular surface, GFP-positive fibers emerged approximately radially from the graft and were broadly distributed among the different quadrants. In progressively deeper optical intervals extending toward the pial surface, however, the relative representation of fibers within the rostral quadrant increased, indicating a gradual transition from a broadly distributed pattern to preferential rostro-directed growth (Fig. 3C).

To compare quantitatively the depth-dependent profiles obtained from all analyzed explants, each preparation was analyzed relative to its own most ventricular interval (z_1_). For each explant, the percentage of GFP signal represented within the anterior quadrant at z_1_ was subtracted from the corresponding values measured at all subsequent optical intervals, such that z_1_ was set to 0. The resulting percentage-point changes in rostral representation were then plotted across the ventricular-to-pial axis for each individual explant, allowing the trajectory of every preparation to be visualized independently. Despite differences in the absolute values and in the magnitude of the response, the individual profiles showed a consistent overall increase in rostral enrichment toward the pial surface (Fig. 3D). Analysis of the original, non-normalized percentages confirmed a significant overall effect of optical depth on rostral axon representation (Friedman test, χ²(3) = 25.56, p < 0.0001; n = 10 explants). These findings indicate that the progressive acquisition of rostro-directed orientation is a reproducible property of donor-derived serotonergic axons as they penetrate the host tissue.

### Donor-derived serotonergic axons follow pre-existing host serotonergic pathways

To determine whether endogenous serotonergic pathways might underlie the depth-dependent rostral polarization of donor-derived axons described above, grafting experiments were repeated using Ai14/Pet1-Cre embryos as host tissue^29,30^. In these preparations, endogenous serotonergic neurons and fibers were selectively labelled with tdTomato, allowing simultaneous visualization of donor-derived GFP-positive axons and host serotonergic pathways. In these preparations, endogenous serotonergic fibers formed a relatively dense and collimated rostrally directed bundle within the ventral hindbrain, providing a well-defined reference pathway against which donor axon behavior could be assessed (Fig. 4A). Consistent with observations in wild-type hosts, donor-derived axons in optical planes closest to the ventricular surface maintained a largely non-oriented distribution. In contrast, in optical planes progressively closer to the pia, a consistent fraction of axons extended alongside endogenous serotonergic bundles and displayed a pronounced rostral orientation (Fig. 4B). In many cases, donor-derived axons could be followed for considerable distances while remaining closely associated with endogenous serotonergic bundles (Fig. 4C). Rather than simply crossing these pathways, GFP-positive fibers frequently adopted parallel trajectories that closely mirrored the orientation of the host serotonergic network (Fig. 4C). These observations raise the possibility that pre-existing serotonergic pathways may provide a permissive physical substrate that contributes to the directional organization of later-growing serotonergic axons.

**Figure 4.**
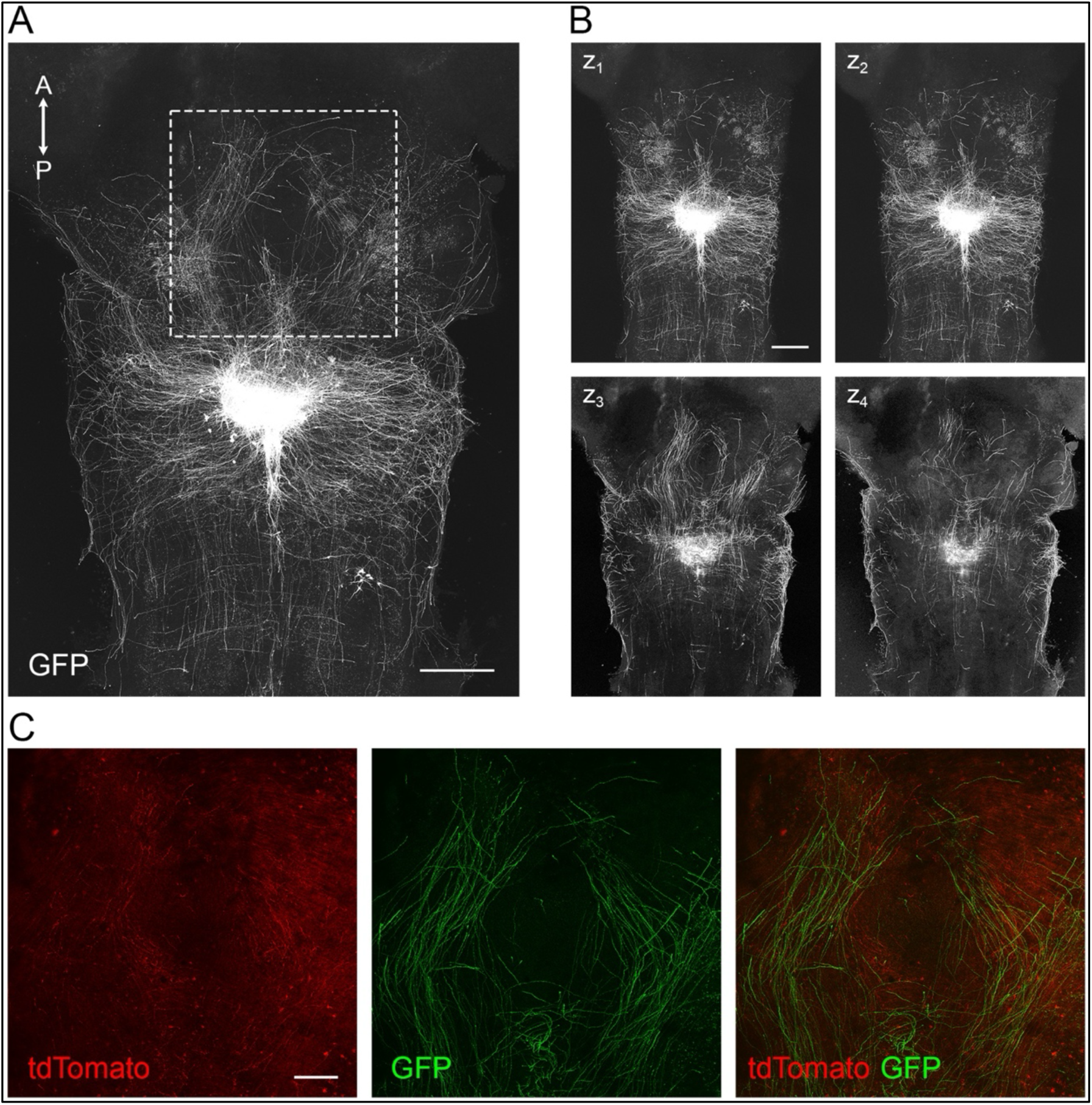
Donor-derived serotonergic axons align with endogenous serotonergic pathways in the developing hindbrain. **(A)** Maximum-intensity projection of GFP-positive axons extending from Tph2-GFP serotonergic neurons homochronically and homotopically grafted onto the ventricular surface of E12.5 Ai14/Pet1-Cre hindbrain flat-mount and maintained 3 days *in vitro*. The dashed box identifies the region shown at higher magnification in C. Anterior-posterior (A-P) orientation is indicated. **(B)** Maximum-intensity projections of four successive optical intervals, z_1_-z_4_, each comprising 10 consecutive confocal sections. Donor-derived axons in the most ventricular intervals, z_1_ and z_2_, display a broadly distributed pattern, whereas in intervals progressively closer to the pia, z_3_ and z_4_, a larger fraction of fibers adopts rostrally-directed trajectories. **(C)** Higher magnification of the boxed region in A showing endogenous tdTomato-positive serotonergic fibers in red, donor-derived GFP-positive axons in green, and the merged channels. Donor-derived axons extend along the trajectories of the endogenous serotonergic pathway. Scale bars: A and B 500 μm; C, 200 μm.

### Donor-derived axons align with host serotonergic pathways across distinct grafting configurations and developmental stages

To further test whether the rostral polarization of donor-derived axons depended specifically on their interaction with endogenous serotonergic pathways, the experimental configuration was reversed by culturing host flat-mount preparations with the ventricular surface facing the membrane and the pial surface exposed upward. In this arrangement, donor explants were positioned directly onto the pial side of the host tissue, so that donor axons encountered endogenous serotonergic streams almost immediately upon entering the host, minimizing the influence of tissue depth as a confounding variable. This configuration also tested whether donor axons would align with and follow the endogenous serotonergic pathway upon contact, rather than producing trajectories with no clear orientation at the top of the preparation (as was the case in the preparation with the ventricular surface up) (Fig. 5A). Donor-derived GFP-positive axons were observed entering endogenous tdTomato-positive bundles and extending alongside them over relatively long distances. At multiple sites, both fiber populations appeared almost superimposed, making their trajectories nearly indistinguishable except for their fluorescent labels (Fig. 5B).

**Figure 5.**
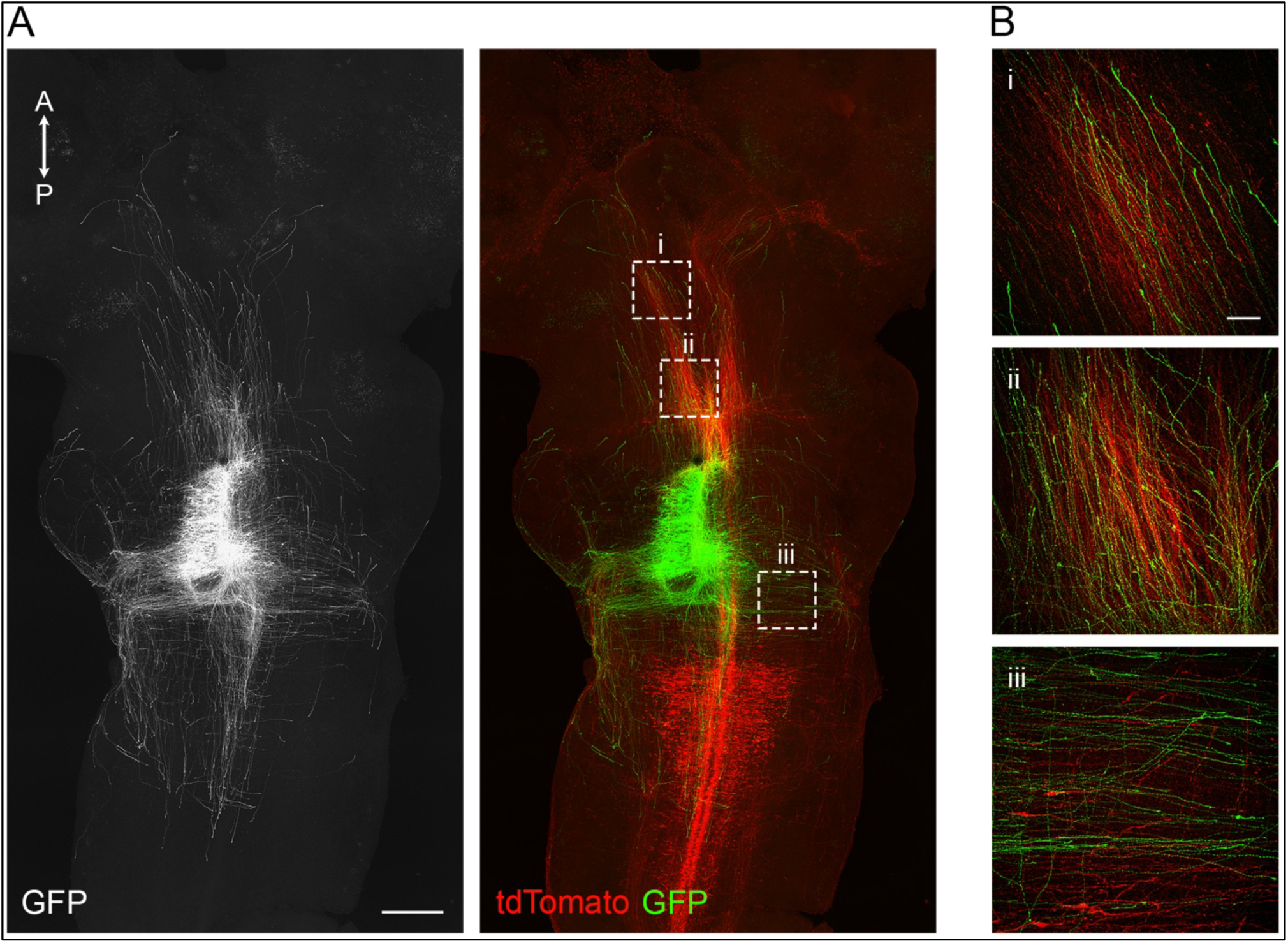
Donor-derived axons follow the local orientation of host serotonergic pathways after pial-side grafting. **(A)** Immunofluorescence showing E12.5 Tph2-GFP serotonergic neurons grafted onto the pial surface of E12.5 Ai14/Pet1-Cre hindbrain flat-mount and maintained 3 days *in vitro* (left). GFP-positive and endogenous tdTomato-positive serotonergic neurons and fibers are shown in the merged image (right). Dashed boxes indicate the regions shown at high magnification in B. Anterior-posterior (A-P) orientation is indicated**. (B)** High magnification of regions i-iii in A. Donor-derived GFP-positive axons maintain a closely parallel course with endogenous tdTomato-positive fibers along rostrally-directed trajectories (**i, ii**) and along a mediolateral component of the host serotonergic pathway (**iii**). Scale bars: A, 500 μm; B, 50 μm.

Because endogenous serotonergic fibers lie immediately beneath the pial surface, this inverted configuration provided improved optical access to the local architecture of the host pathway compared to grafts with the ventricular surface facing upward (in which most serotonergic axons were located deeper in the host tissue). This revealed that the endogenous projection was not directed exclusively rostrally but also contained prominent mediolateral trajectories (see panel iii in Fig. 5B). Notably, donor-derived fibers were scarce both in deeper optical planes extending beyond the endogenous serotonergic pathway toward the ventricular side and in pial surface regions lacking host serotonergic fibers. This distribution argues against a simple random spread of donor axons throughout the host tissue and supports a close spatial association between donor and host serotonergic axons. The rostral polarization observed in this pial grafting configuration was consistent with that previously described for grafts placed on the ventricular surface, supporting the idea that directional organization reflects an interaction with endogenous serotonergic pathways rather than a nonspecific effect of tissue depth.

To further challenge this interpretation, we repeated the experiment using E10.5 host preparations. At this developmental stage, serotonergic neurons are only beginning to differentiate and have not yet established the mature ventral serotonergic fiber bundles characteristic of later embryonic stages. Importantly, because the E10.5 neural tube is considerably thinner than at E12.5, donor explants placed on the ventricular surface remained in close proximity to the earliest differentiating serotonergic neurons, approximating the spatial relationship achieved in the E12.5 pial-side grafting configuration. Under these conditions, donor-derived axons should encounter the earliest pioneer serotonergic fibers almost immediately upon emerging from the graft. Remarkably, despite the immaturity of the host serotonergic system, donor-derived axons displayed a pronounced rostral orientation (Fig. 6), suggesting that even the earliest pioneer serotonergic fibers present at this stage may provide orientational information to later-growing axons.

**Figure 6.**
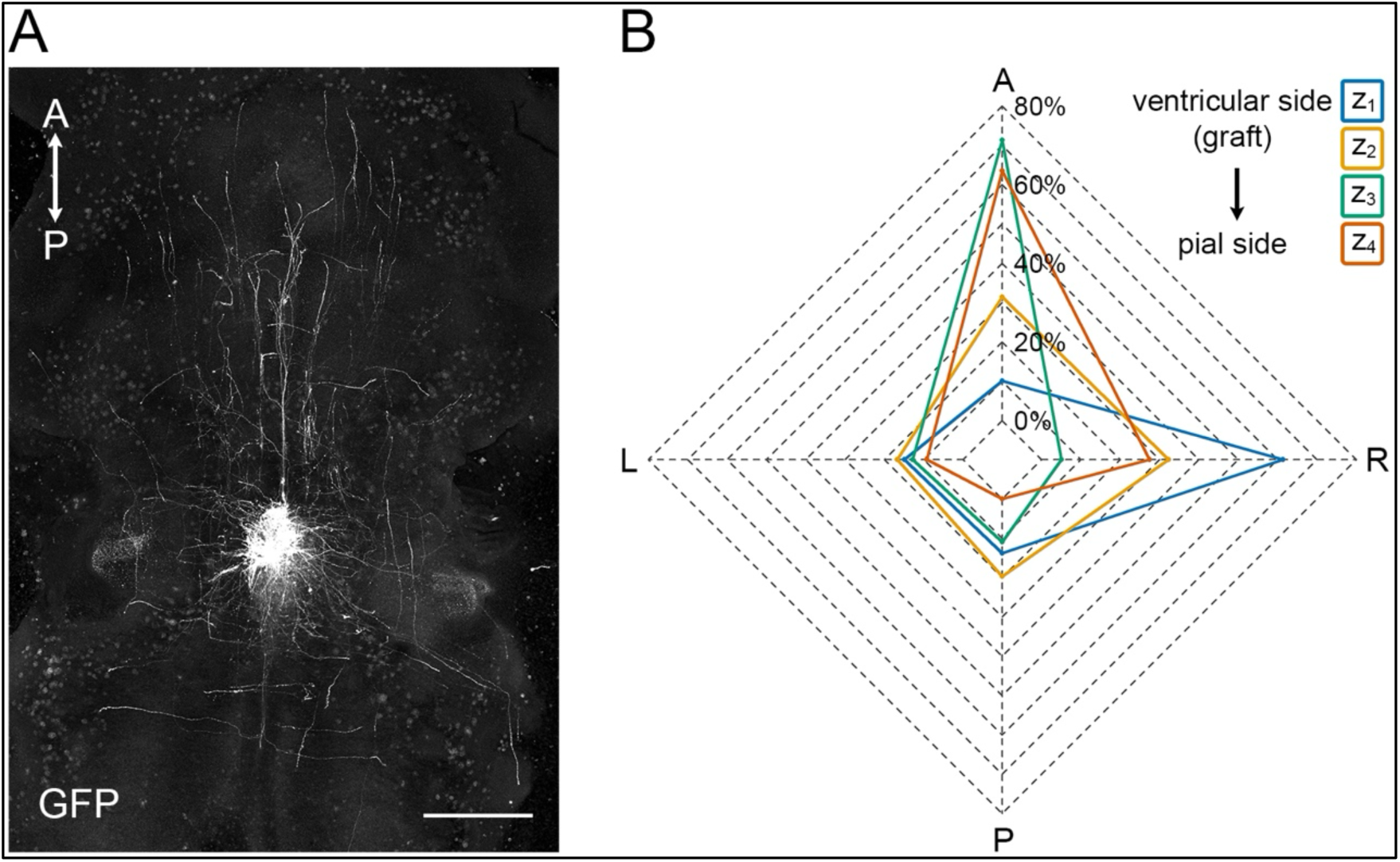
Donor-derived serotonergic axons acquire rostro-directed orientation in E10.5 host tissue. **(A)** Maximum-intensity projection of GFP-positive axons extending from an E12.5 Tph2-GFP rostral serotonergic neurons grafted onto the ventricular surface of E10.5 hindbrain flat-mount and maintained 3 days *in vitro*. A clear anterior bias in the distribution of donor-derived fibers is already evident in the full z-stack projection. Anterior-posterior (A-P) orientation is indicated**. (B)** Radar plot showing the percentage contribution of the anterior, posterior, left, and right quadrants to the GFP relative optical density measured in four successive optical intervals, z_1_-z_4_, extending from the ventricular surface toward the pial surface. The intervals closer to the pia display pronounced enrichment of donor-derived fibers within the anterior quadrant. A, anterior; P, posterior; L, left; R, right. Scale bar: A, 500 μm.

These observations, taken together across two distinct grafting configurations and two developmental stages, indicate that the directional organization of donor-derived serotonergic axons is a robust phenomenon that emerges from early contact with endogenous serotonergic pathways. The near-complete restriction of donor-derived fibers to optical planes containing the host serotonergic stream further suggests that these pathways may constrain the trajectories of newly entering axons. This is consistent with the idea that pre-existing serotonergic fibers act as permissive and potentially instructive substrates for the navigation of later-growing serotonergic axons.

### Donor-derived axons adapt their trajectories to heterotopic serotonergic pathways

The experiments described above showed that donor-derived serotonergic axons align with endogenous serotonergic fibers and adopt the local orientation of the host pathway. We next asked whether this behavior was restricted to pathways normally encountered by rostral raphe axons or could also occur within a heterotopic serotonergic environment. To address this, rostral raphe explants from rhombomeres 1-2 were transplanted at the level of rhombomeres 6-7, within the territory of the developing caudal raphe, whose neurons display a distinct molecular identity and project predominantly toward the spinal cord^31,32^.

Donor-derived axons extended along the serotonergic pathways present within the caudal raphe territory. Most fibers followed the caudally directed stream toward the spinal cord or adopted mediolateral and rostral trajectories (Fig. 7A). Overall, the distribution of donor-derived axons closely reflected the organization of the local serotonergic network, indicating that rostral raphe axons can adapt their trajectories to pathways encountered within a heterotopic host environment (Fig. 7B).

**Figure 7.**
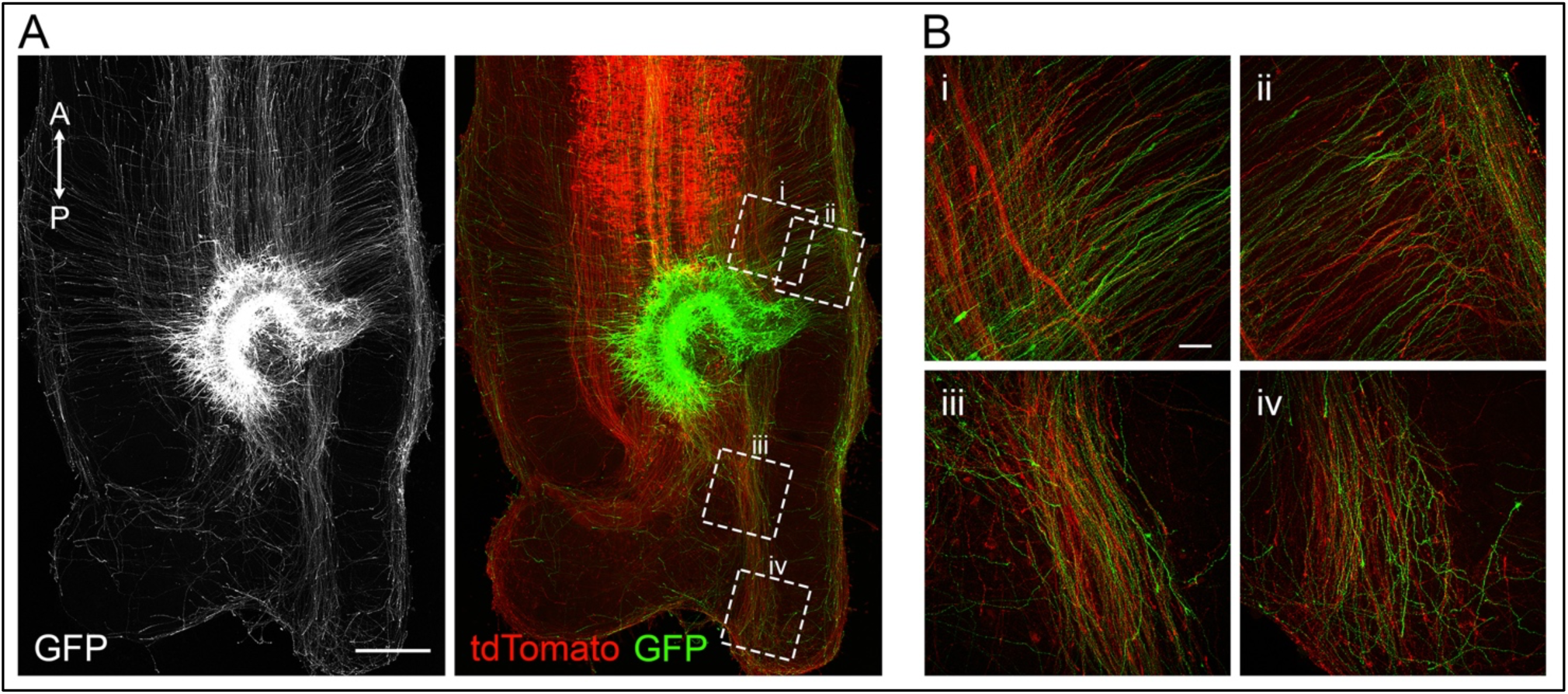
Donor axons adapt their trajectories to a heterotopic caudal serotonergic environment. **(A)** Immunofluorescent images showing E12.5 donor GFP-positive neurons from rhombomeres 1-2 heterotopically transplanted into the caudal hindbrain territory (rhombomeres 6-7) of Ai14/Pet1-Cre embryonic hindbrain flat-mount and maintained 3 days *in vitro* (left). GFP-positive donor-derived and endogenous tdTomato-positive serotonergic neurons are shown in the merged image (right). Dashed boxes identify the regions shown in B. Anterior-posterior (A-P) orientation is indicated**. (B)** High magnification of regions i-iv in A. Donor-derived GFP-positive axons extend in close parallel association with local tdTomato-positive serotonergic pathways and adopt the range of trajectories present within the host caudal raphe territory, including longitudinal, mediolateral, and caudally directed components. Scale bars: A, 400 μm; B, 50 μm.

These findings indicate that rostral serotonergic axons retain substantial navigational plasticity and are capable of engaging with serotonergic pathways that they would not normally encounter in their native territory. Although the present experiment does not distinguish whether this behavior is driven primarily by direct interactions with host fibers or by molecular features of the surrounding caudal environment, it shows that donor axons are not irrevocably committed to a fixed rostro-directed program but remain responsive to the organization of a heterotopic serotonergic system.

### Embryonic serotonergic axons retain robust growth capacity and respond to tissue architecture in adult brain slices

To determine whether embryonic serotonergic neurons retain their growth potential and environmental responsiveness within a mature neural context, rostral raphe explants from E12.5 Tph2-GFP embryos were transplanted onto adult organotypic brain slices derived from wild-type animals and maintained *in vitro* for five days (5 DIV) under the same culture conditions (Fig. 8A). Because the adult brain represents a substantially more growth-restrictive environment than embryonic tissue, the ability of early serotonergic neurons to survive and extend axons under these conditions was not self-evident. Donor-derived serotonergic neurons survived efficiently on the adult host tissue and extended abundant GFP-positive axons throughout the surrounding parenchyma spreading over large portion of the host slices and, demonstrating that embryonic serotonergic neurons retain considerable intrinsic growth capacity even when placed in a mature cellular environment (Fig. 8B,C).

**Figure 8.**
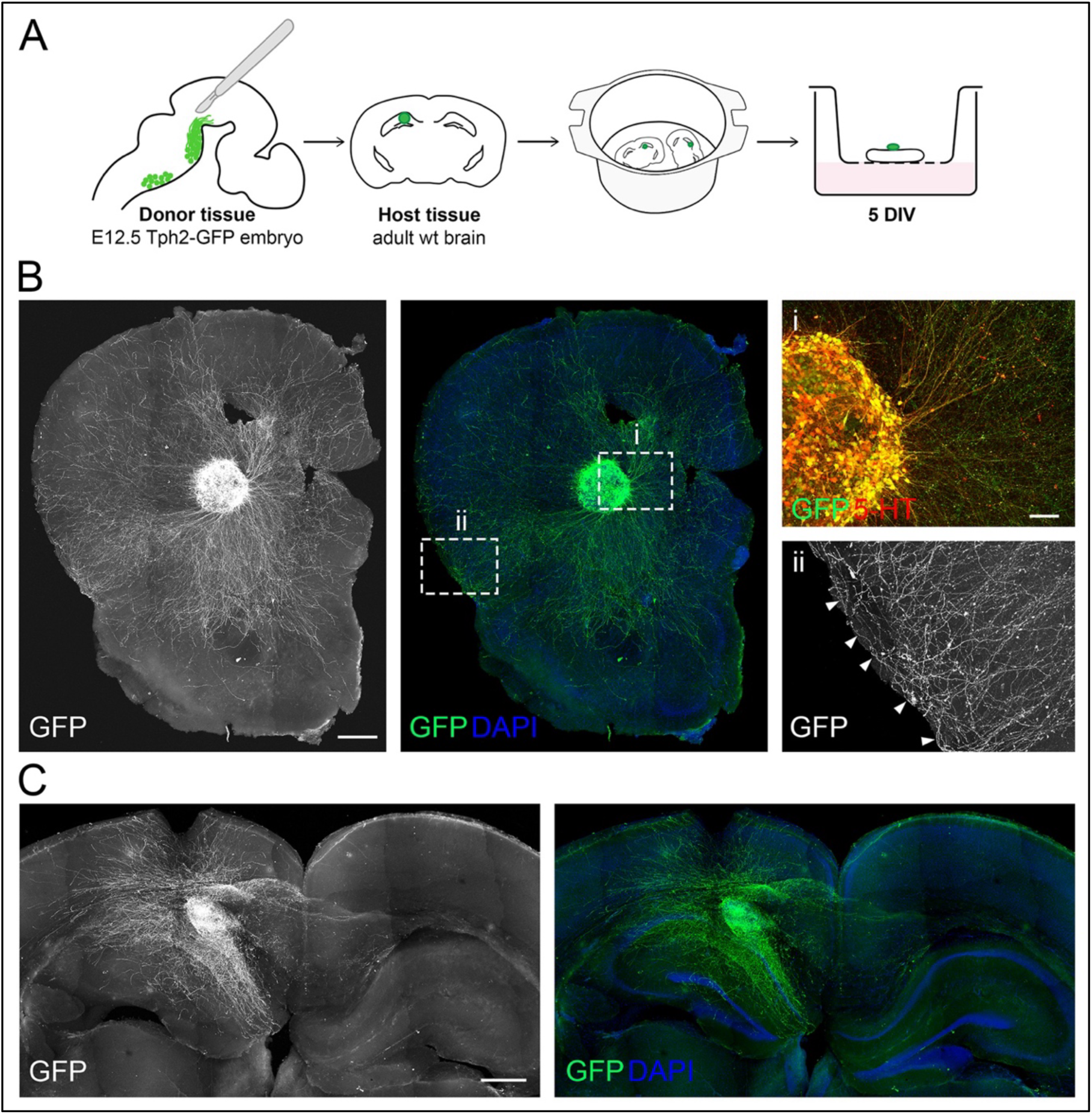
Embryonic serotonergic axons retain growth competence and respond to adult brain tissue architecture. **(A)** Schematic overview of the heterochronic transplantation procedure. Rostral raphe serotonergic neurons were dissected from E12.5 Tph2-GFP embryos, grafted onto organotypic slices of adult wild-type mouse brain, and maintained 5 days *in vitro*. **(B)** Representative adult coronal brain slice at the level of the rostral forebrain containing an E12.5 Tph2-GFP donor explant. The GFP channel is shown in grayscale (left) and together with DAPI counterstaining (right). Donor-derived GFP-positive axons extend throughout the surrounding adult host tissue. Dashed boxes indicate the regions shown at higher magnification in panels i and ii. **(i)** Double immunolabeling for GFP and 5-HT confirms the serotonergic identity of donor-derived cells and fibers. **(ii)** GFP-positive axons reaching the cortical surface reorient and extend tangentially along the pial boundary and cortical layer I (arrowheads). **(C)** Representative adult coronal brain slice containing the hippocampal formation with an E12.5 Tph2-GFP donor explant. The GFP channel is shown in grayscale (left) and together with DAPI counterstaining (right). Donor-derived axons extend over long distances within the host tissue following hippocampal laminar boundaries. 5-HT, serotonin; DIV, days *in vitro*. Scale bars: B, 500 μm; Bi-ii, 100 μm; C, 500 μm.

Unlike the polarized growth observed in embryonic hindbrain preparations, donor-derived axons on adult slices did not display a consistent overall directional bias. Axons extended over long distances, often following tortuous paths and giving rise to numerous collateral branches. Nevertheless, their trajectories did not appear completely random in that growing axons frequently adjusted their course in relation to the local organization of the host tissue rather than crossing the parenchyma indiscriminately. In cortical regions, fibers reaching the pial surface reoriented and extended tangentially along layer I, closely matching the orientation of endogenous serotonergic fibers at this depth (see panel ii in Fig. 8B). In hippocampal slices, donor-derived axons preferentially followed laminar boundaries instead of traversing the tissue directly (Fig. 8C). These observations extend the navigational plasticity revealed by the embryonic grafting experiments and indicate that developing serotonergic axons remain responsive to structural features of the host environment even in adult tissue. Although the relevant substrates are not identified here, the tendency of donor-derived axons to align with cortical and hippocampal boundaries is consistent with a broader capacity to exploit pre-existing tissue architecture during growth.

Together, these findings indicate that developing serotonergic axons are strongly influenced by pre-existing serotonergic pathways during navigation, consistent with the classical hypothesis of epiphytic guidance proposed decades ago based on neuroanatomical observations^1–3^ but has remained untested experimentally. By allowing donor-derived axons to be observed in direct relation to genetically labelled endogenous serotonergic fibers, our transplantation paradigm provides experimental support for this model and extends it toward a pioneer-follower framework similar to that described in *Drosophila*, zebrafish and *C. elegans*^22–24^. Rather than emerging already committed to a fixed trajectory and navigating following diffusible cues, later-growing serotonergic axons appear to be able to use pre-existing serotonergic bundles as physical substrates for their own growth, progressively acquiring directional organization as they approach the host pathway. The heterotopic transplantation experiments further support this interpretation. Rather than simply maintaining the predominantly rostro-directed organization characteristic of their native territory, donor axons adopted the range of trajectories present within the caudal host network.

This indicates that regional molecular identity alone does not rigidly determine axon trajectory *a priori*, and that donor axons retain substantial navigational plasticity, although the present experiments cannot fully distinguish whether this behavior reflects a direct response to endogenous fibers themselves or to molecular cues distributed along their trajectory. Nonetheless, the presence of directional organization already at E10.5, before mature longitudinal bundles have formed, suggests that even the earliest pioneer fibers can provide orientational information to later-growing axons. These observations support a hierarchical model in which initial trajectories, likely established through classical guidance mechanisms, subsequently become substrates for the progressive, self-generated expansion of the serotonergic network. This view is also compatible with previous explant-based co-culture studies showing that developing dorsal raphe serotonergic axons can respond to short-range, tissue-associated guidance cues *in vitro*^33^. In the model proposed here, such progressive self-scaffolding would considerably reduce the positional information that must be specified for each individual axon, allowing an extensive projection system to emerge from comparatively simple local interaction rules. This mechanism does not, however, explain the striking regional diversity of mature serotonergic innervation, which varies markedly in axon caliber, branching pattern, varicosity density, and reliance on wiring versus volume transmission across brain territories^1,34–37^. We speculate that, while fasciculation with pioneer bundles guides axons toward broad target territories, the finer features of terminal arborization and the local balance between transmission modes are established only after arrival, by interpreting region-specific molecular cues. This implies that raphe neurons may possess a broad molecular competence to respond appropriately to local signals, rather than a fully pre-specified, target-selective program.

Finally, embryonic serotonergic axons transplanted onto adult brain slices retained a comparable ability to interpret local tissue architecture, extending along cortical layer I and hippocampal laminar boundaries. Fiber accumulation in these regions may be further supported by the stochastic properties of serotonergic fibers as they interact with physical tissue borders^38,39^. Although the serotonergic system is known to retain a high degree of structural plasticity well into adulthood^40,41^, the mature brain likely no longer contains actively growing pioneer serotonergic pathways; nonetheless, embryonic serotonergic axons remained highly responsive to the structural organization of the host tissue rather than extending indiscriminately through the parenchyma.

This indicates that substrate-dependent navigation is a general property of serotonergic axons that is not restricted to the embryonic period. One possibility is that this robust growth capacity is facilitated by the widespread distribution of serotonergic fibers throughout the adult brain, which may provide abundant pre-existing axonal substrates and structural routes that can be exploited by newly extending serotonergic axons. Interestingly, such robust growth is not necessarily a general property of embryonic monoaminergic neurons placed in organotypic tissue. In a recent study, grafted embryonic A9 dopaminergic neurons displayed only limited extension of TH-positive neurites beyond the graft^27^. Although differences in neuronal identity and experimental design preclude a direct comparison, these observations raise the possibility that serotonergic axons possess an intrinsic broader capacity to exploit pre-existing tissue architecture for growth and navigation.

Several questions remain open. The present experiments do not distinguish whether donor axons interact directly with serotonergic membranes, with axon-associated extracellular components or with neighboring cellular structures organized around the host pathway. Nor do they establish whether host serotonergic fibers are strictly required for donor-axon orientation. Selective disruption of the endogenous pathway, together with live imaging of growth-cone behavior, will be needed to resolve the underlying mechanism and to determine whether contact promotes turning, stabilization, fasciculation or preferential elongation.

Overall, our findings support a model in which serotonergic pathway assembly emerges from the interaction between intrinsic growth competence, classical tissue-derived guidance, and progressive use of pre-existing axonal substrates. By allowing early projections to guide later-growing fibers, this mechanism offers a plausible explanation for how a relatively small population of raphe neurons can generate a widespread yet spatially organized neuromodulatory system.

## Methods

### Animals

Mice were housed in standard Plexiglas cages at a constant temperature of 22 ± 1 °C under a 12-h light/12-h dark cycle, with food and water available *ad libitum*. All experimental procedures were conducted in accordance with the guidelines of the Ethics Committee of the University of Pisa and were approved by the Veterinary Department of the Italian Ministry of Health (AC179.N.HQV and AC179.N.CCL).

### Organotypic cultures

Embryos from timed-pregnant C57BL/6J mice were collected at embryonic day (E) 10.5 or E12.5, as described in Pasqualetti et al.^42^, and immediately transferred to ice-cold artificial cerebrospinal fluid (aCSF) containing 1 mM NaH₂PO₄, 4 mM glucose, 24 mM NaHCO₃, 124 mM NaCl, 5 mM KCl, 2 mM CaCl₂, and 1 mM MgCl₂ (Merck). Brains from wild-type and Ai14/Pet1-Cre embryos^29,30^ were used as host tissue for transplantation experiments. Brains were dissected under a Nikon SMZ18 stereomicroscope equipped with fluorescence illumination and carefully isolated from surrounding tissues and meninges. To preserve the overall rostrocaudal organization of the developing tissue, whole brains were opened along the dorsal midline and flattened to obtain flat-mount preparations. Samples were positioned on polyethylene terephthalate (PET) membrane cell-culture inserts with 1-μm pores (Corning), with the ventral surface facing either upward or downward. The inserts were placed in multiwell culture plates containing DMEM/F12 (Thermo Fisher Scientific) supplemented with 30% horse serum (Merck), 36 mM glucose (Merck), and 1% penicillin/streptomycin (Merck).

For adult organotypic cultures, wild-type mice were euthanized by cervical dislocation, and their brains were rapidly dissected, embedded in 4% low-melting point agarose prepared in aCSF, and sliced coronally at a thickness of 200 μm in ice-cold aCSF using a vibratome (Leica Microsystems). Slices were collected, transferred onto PET membrane cell-culture inserts, and maintained under the same culture conditions described above.

Donor tissue was obtained from the rostral raphe region of heterozygous Tph2-GFP embryos^28^. After dissection, the hindbrain was prepared as a flat-mount, and the rostral portion of one fluorescent serotonergic stripe, corresponding approximately to the ventral half of rhombomeres 1 and 2, was isolated using an ophthalmic microsurgical scalpel. A medial longitudinal cut separated the two sides of the hindbrain, while transverse and lateral cuts delineated the r1-r2 donor fragment containing the fluorescent serotonergic neurons. Grafts were positioned on flat-mount preparations or organotypic slices using glass capillaries and fine tungsten needles. Cultures were maintained at 37 °C in a humidified atmosphere containing 5% CO₂ for 3 days for embryonic preparations or 5 days for adult brain-slice cultures.

### Immunohistochemistry

Organotypic cultures were fixed overnight in 4% paraformaldehyde (Merck) at +4 °C. For immunohistochemical processing, the membrane surrounding each preparation was carefully trimmed, and samples were processed as free-floating preparations. Fluorescence immunohistochemistry was performed as previously described^43^. Briefly, samples were incubated for 1 h at room temperature in a blocking solution containing PBS, 0.5% Triton X-100, and 5% lamb serum (Merck), followed by overnight incubation at +4 °C with one of the following combinations of primary antibodies: rabbit anti-GFP (1:2,000; Thermo Fisher Scientific) and chicken anti-RFP (1:500; Synaptic Systems), or chicken anti-GFP (1:1,000; Abcam) and rabbit anti-5-HT (1:500; Merck). The following day, samples were washed three times in PBS containing 0.5% Triton X-100 and incubated overnight at +4 °C with the corresponding combinations of secondary antibodies: Alexa Fluor 488 donkey anti-rabbit and Alexa Fluor 555 donkey anti-chicken, or Alexa Fluor 488 donkey anti-chicken and Alexa Fluor 594 donkey anti-rabbit (all diluted 1:500; Thermo Fisher Scientific). Samples were then washed three times in PBS containing 0.5% Triton X-100, counterstained with DAPI (1:1,000; Thermo Fisher Scientific), and mounted using Fluor Guard mounting medium (Histo-Line).

### Confocal imaging and relative optical density analysis

Fluorescence images were acquired using a Nikon AX confocal laser-scanning microscope equipped with a 10× objective with a numerical aperture of 0.45. Large-area tile-scan acquisitions were performed to image the entire embryonic flat-mount preparation or adult organotypic brain slice. Images were acquired at a resolution of 1024 × 1024 pixels with a z-step of 2 μm. Additional high-magnification images were acquired using 20× and 40× objectives for qualitative visualization of axonal morphology and donor-host fiber interactions presented in the figures; these images were not used for quantitative analysis.

Serotonergic fiber distribution at different depths within the host tissue was quantified by relative optical density (ROD) analysis, as previously described^44^. Briefly, z-stacks were divided into groups of 10 consecutive optical sections, and a maximum-intensity projection was generated for each group. Based on the preserved rostrocaudal organization of the preparation, the host tissue surrounding the graft was divided into four quadrants corresponding to the anterior, posterior, left, and right directions. The edge of each donor explant was manually delineated to account for its irregular shape, and the explant core was excluded from the analysis. ROD measurements were then performed on GFP-positive serotonergic fibers within the remaining area of each quadrant. Background optical density (OD) was measured in regions within the same analyzed area that lacked GFP-positive serotonergic fibers, and ROD was calculated by subtracting the background OD from the measured signal. ROD values were calculated independently for each quadrant across successive groups of optical sections using Fiji/ImageJ software. This approach enabled the depth-dependent analysis of axonal distribution and directional growth of graft-derived serotonergic fibers within the host tissue. Radar plots representing quadrant-specific ROD measurements across different optical depths were generated in R (version 4.4.0) using the fmsb (version 0.7.6) and RColorBrewer (version 1.1-3) packages. For visualization of depth-dependent changes in rostro-directed growth, the percentage contribution of the rostral quadrant to the total GFP ROD was plotted against optical depth for each individual explant using the ggplot2 package (version 3.5.1). To facilitate comparison of depth-dependent trajectories among explants, the value measured in the most ventricular optical interval was subtracted from the values measured in all intervals of the same preparation, such that the most ventricular interval was set to zero. Values therefore represent percentage-point changes relative to the most ventricular interval. Statistical analyses were performed on the original, non-normalized percentages.

## Statistical analysis

Statistical analyses were performed in R (version 4.4.0). Because the data were not normally distributed, nonparametric statistical tests were used. Differences in the percentage contribution of the rostral quadrant across successive optical intervals were evaluated using a Friedman test for repeated measures. Statistical significance was set at *p* < 0.05.

## Author contributions

M.Pi. and M.Pa. designed and performed research, and wrote the paper; M.Pi., S.N., S.M., G.G., N.B. and M.Pa. analyzed data; SJ contributed to the theoretical hypothesis; S.N., S.M., G.G., S.J., and N.B. contributed to the preparation of the manuscript.

## Acknowledgments

We thank C. Valente for excellent technical assistance and all members of our laboratory for valuable discussions and comments on the manuscript.

